# Interactive effects of aging and aerobic capacity on energy metabolism-related metabolites of serum, skeletal muscle, and white adipose tissue

**DOI:** 10.1101/2020.07.16.207746

**Authors:** Haihui Zhuang, Sira Karvinen, Xiaobo Zhang, Xiaowei Ojanen, Timo Törmakangas, Vidya Velagapudi, Markku Alen, Steven L. Britton, Lauren G. Koch, Heikki Kainulainen, Sulin Cheng, Petri Wiklund

**Author notes:** Equal contribution (Haihui Zhuang, Sira Karvinen). Equal contribution (Heikki Kainulainen, Sulin Cheng, Petri Wiklund).

## Abstract

Aerobic capacity is a strong predictor of longevity. With aging, aerobic capacity decreases concomitantly with changes in whole body metabolism leading to increased disease risk. To address the role of aerobic capacity, aging and their interaction on metabolism, we utilized rat models of low and high intrinsic aerobic capacity (LCRs/HCRs) and assessed the metabolomics of serum, muscle, and white adipose tissue (WAT). We compared LCRs and HCRs at two time points: Young rats were sacrificed at 9 months, and old rats were sacrificed at 21 months. Targeted and semi-quantitative metabolomics analysis was performed on ultra-pressure Liquid Chromatography Tandem Mass Spectrometry (UPLC-MS) platform. Interaction of aerobic capacity and aging was studied via regression analysis. Our results showed at young age, metabolites linked to amino acid metabolism differed in serum and muscle with aerobic capacity, whereas no difference were observed in WAT. In aged animals, most prominent changes in metabolites occurred in WAT. Aerobic capacity and aging interactively affected seven metabolites linked to energy metabolism. Our results support previous findings that high aerobic capacity is associated with more efficient amino acid metabolism in muscle. While impaired branched chain amino acids (BCAAs) and fatty acid metabolism in the muscle may associate to the high risk of metabolic disorders and shorter lifespan previously observed in LCRs. The interactive effects of aging and aerobic capacity on energy metabolism-related metabolites were largely driven by HCRs, reflecting the importance of inherited aerobic capacity in the aging process. Our results highlight that dysfunctional mitochondrial β-oxidation in WAT may be one key mechanism related to aging.

## INTRODUCTION

Epidemiological studies have indicated that low aerobic capacity, indexed as maximal oxygen uptake (VO_2max_), is a major risk factor for cardio-metabolic diseases [1], whereas high aerobic capacity is associated with a more favorable cardio-metabolic health profile [2, 3], even in the presence of excess adiposity [4]. These observations are complemented by multiple randomized controlled trials, which have shown that exercise training significantly improves cardiorespiratory fitness and adiposity-related cardio-metabolic biomarkers [5]. Furthermore, aerobic capacity has been shown to be a strong independent predictor of cardiovascular and all-cause mortality in many population cohorts [6]. These studies suggest there is a fundamental connection between impaired aerobic metabolism, complex disease risk, and reduced longevity.

Epidemiological studies have also shown that aerobic capacity and physical activity level decrease with age [7, 8] concomitantly with quantitative and structural changes in body composition [9]. These changes are associated with alterations in whole body metabolism, insulin resistance, low-grade inflammation and dyslipidemia, which contribute to the development of cardio-metabolic disease with age [10, 11]. However, it is not well understood to what extent the aging-related changes in metabolic profiles are attributable to either intrinsic aerobic capacity or decreased physical activity.

To enable the investigation of intrinsic aerobic endurance capacity on metabolic profile, Koch and Britton [12] developed heterogeneous rat strains by artificial selection for low and high inborn exercise capacity. This rat model of low capacity runners (LCRs) and high capacity runners (HCRs) prospectively tests the association between aerobic exercise capacity and survivability [13]. The phenotype of HCRs is coincident with a host of health benefits [14], including a 28–40% increased lifespan [15]. A recent study in HCRs and LCRs showed that an enhancement of aerobic capacity could mitigate some of the changes in the plasma metabolic profile, which were associated with aging [16]. Concomitantly with higher aerobic capacity, muscle mitochondrial function and oxidative energy metabolism are enhanced in HCRs compared with LCRs [17, 18]. In addition to skeletal muscle, white adipose tissue (WAT) shares an important role in the regulation of whole body metabolism [19] and WAT-to-muscle communication is critical in overall metabolic health [20].

Most previous studies have examined the effects of aerobic capacity on metabolism only in one tissue [21, 22] and using only young animals (3-4 months of age) [17]. Thus, possible interactive effects of aging and aerobic capacity on tissue metabolism are yet to be resolved. Here, we assessed metabolic profiles in serum, muscle and WAT in both young (9 months of age) and old (21 months of age) HCRs and LCRs groups by using targeted and semi-quantitative metabolomics analysis which was performed on the triple quadrupole tandem mass spectrometry coupled to ultra-pressure liquid chromatography (UPLC-MS) platform. We propose to define the most prominent differences in the metabolic profiles of LCRs and HCRs in skeletal muscle [17, 18, 23, 24]. Also, we expect to observe aging-related differences in metabolites particularly in LCRs, as LCRs are known to have higher metabolic disease risk and shorter lifespan [13, 14].

## MATERIALS AND METHODS

### Animal model

The HCR/LCR rat model was derived from a genetically heterogeneous founder population (N: NIH stock) and bred with two-way artificial selection [12]. Animals were evaluated at 11 weeks of age for maximal running capacity at the University of Michigan (Ann Arbor, Michigan, USA) with a speed-ramped treadmill running test (15 slope, initial velocity of 10 m/min, increased 1 m/min every 2 min). In this study, 34 female rats (16 HCRs and 18 LCRs) from generations 23-27 of selection were used. All rats were kept in an environmentally controlled facility with light/ dark cycle of 12/12 hours and had free access to food and tap water (R36, Labfor, Stockholm, Sweden).

### Testing procedure

After arriving to Finland, rats were tested for maximal running capacity at the age of 9 months with the same speed-ramped running test as described previously [18]. Subsequently, both HCRs and LCRs were divided into weight-and maximal running capacity-matched sub-groups: HCR-Y or LCR-Y (young, n = 10 in each group), and HCR-O (old, n = 6) or LCR-O (old, n = 8). After maximal aerobic capacity testing, HCR-Y and LCR-Y were weighed, sacrificed and hindlimb skeletal muscles, WAT and blood samples were collected. HCR-O and LCR-O were continued to be housed individually in a standard cage until the age of 21 months. Maximal aerobic capacity was assessed again at the age of 21 months. After testing, animals were weighed, sacrificed and skeletal muscle, WAT and serum samples were harvested.

### Tissue collection

Soleus, extensor digitorum longus (EDL), plantaris, gastrocnemius, quadriceps femoris muscle, adipose tissue around the ovaries, visceral adipose tissue and retroperitoneal adipose tissue were excised and weighed. Tissue samples were snap frozen in liquid nitrogen then stored in −80□ until analyses. Gastrocnemius, retroperitoneal adipose tissue and serum were used for metabolomics analyses. Skeletal muscle mass was calculated as the sum weight of soleus, EDL, plantaris, gastrocnemius and quadriceps femoris.

### Metabolomics analyses

Targeted and semi-quantitative metabolomics analyses were performed on a Waters Xevo TQ-S triple quadrupole tandem mass spectrometer coupled to ultra-pressure liquid chromatography (UPLC-MS) platform using the previously published protocol in FIMM (Institute for Molecular Medicine Finland) [25]. Briefly, metabolites were extracted from 100 µl serum, 20 mg muscle and 20mg WAT samples respectively using protein precipitation by adding acetonitrile +1% formic acid. The collected extracts were dispensed in Ostro 96-well plates (Waters Corporation, Milford, USA) and filtered by applying a vacuum at a delta pressure of 300–400 mbar for 2.5□min on robot’s vacuum station. Filtered sample extract (5□µL) was injected in an Acquity UPLC-system coupled to a Xevo TQ-S triple quadrupole mass spectrometer (Waters Corporation, Milford, MA, USA) which was operated in both positive and negative polarities with switching time of 20 milliseconds. Multiple Reaction Monitoring (MRM) acquisition mode was selected for the quantification of metabolites. MassLynx 4.1 software was used for data acquisition, data handling and instrument control. Data processing was done using TargetLynx 4.1 software. 89 metabolites in serum, 71 in muscle and 71 in WAT were identified in whole samples.

### Data analysis

The descriptive characteristics of rats were analyzed by using IBM SPSS Statistics 24.0 (SPSS, Chicago, USA). Data were checked for normality with the Shapiro-Wilks test. As most of the variables were not normally distributed, nonparametric tests were chosen for group comparisons, and statistical significance was set at *p* < 0.05.

All metabolomics data were normalized by the sum and followed by log transformation. Thereafter, data was exported to SIMCA-P+ 14.0 software (Umetrics, Sweden) for multivariate analysis. The data were UV scaled, and an orthogonal partial least-squares-discriminant analysis (OPLS-DA) statistical models were used to compare differences between groups [26]. Then a permutation test (n = 200) was used to validate the model and to avoid over-fitting. Furthermore, the variable influence of projection (VIP) values of the OPLS-DA models were used to identify the most influential metabolites contributing to group separation. Metabolites with VIP□≥□1 and p value < 0.05 were considered to differ between groups.

We used linear regression models to assess the differential association of each metabolite separately on aerobic capacity while using age and HCR-Y, HCR-O, LCR-Y and LCR-O groups as design factors. From these models, we report the interaction unstandardized coefficients of the metabolites with the design factors. Where necessary, we used standard variable transformations to achieve normal distribution of the residuals from the regression models and to control for outliers in predictor variables.

### Pathway analysis

Metaboanalyst (v3.5) was used to identify metabolic pathways for mapping altered metabolites to their corresponding pathway.[27, 28]. This web-based tool relies on the knowledgebase of Kyoto Encyclopedia of Genes and Genomes (KEGG) metabolic pathway.

## RESULTS

### Body mass, running capacity and skeletal muscle mass/body mass ratio

HCRs had lower body mass and higher maximal running speed, a correlate to VO_2max_ than LCRs at both 9 months (p < 0.001, p < 0.001) and 21 months of age (p = 0.002, p = 0.006) (**Fig. 1a, 1a**). The maximal running speed was 25.5% lower in 21 months old HCRs compared with 9 month old HCRs (p < 0.001) whereas in LCRs, no significant difference in running speed with age was observed (p = 0.360). Hindlimb skeletal muscle mass to body mass ratio was lower in both HCRs and LCRs when comparing 21 months old rats to 9 months old (**Fig. 1b**).

**Figure. 1.**
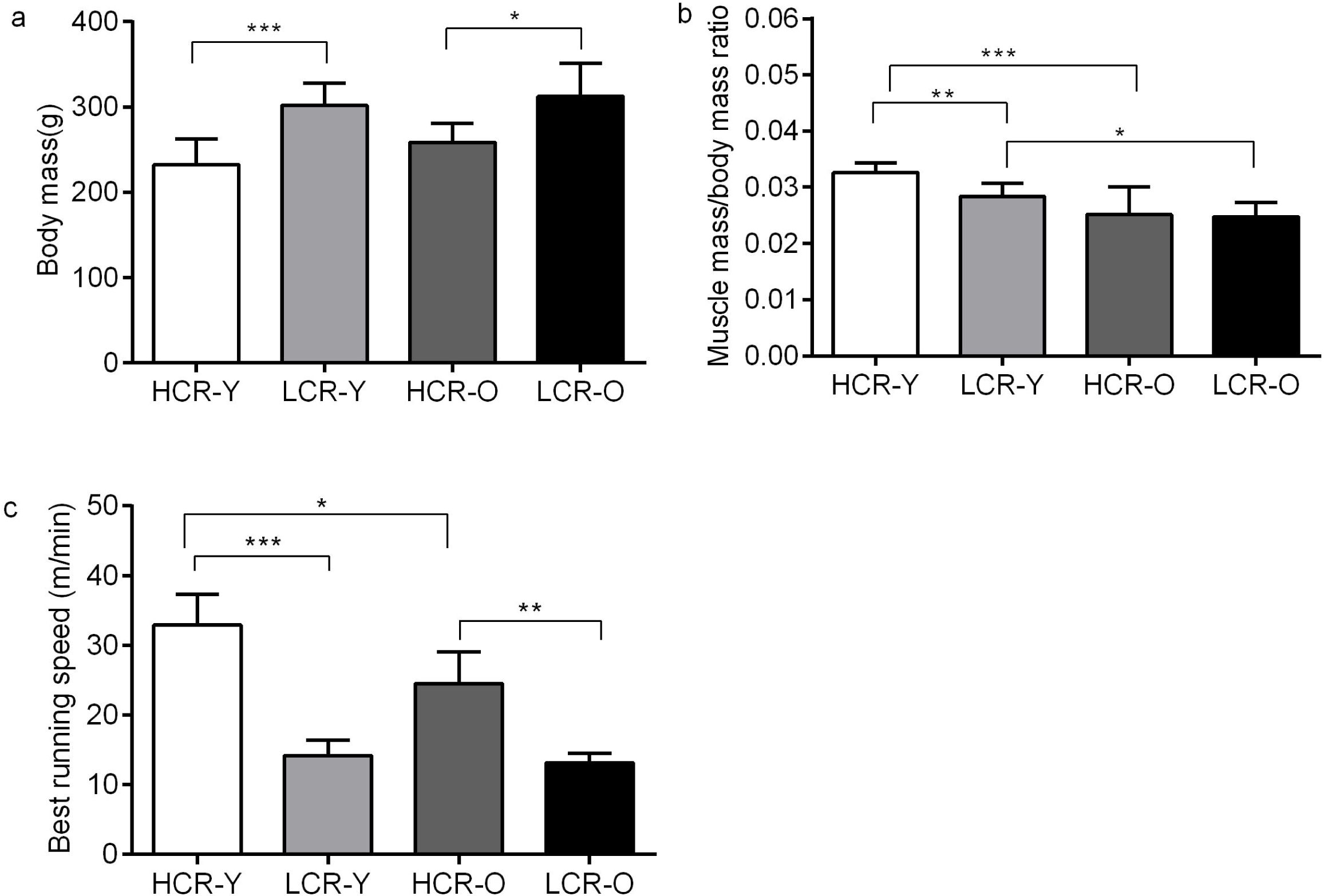
The effects of aerobic capacity and aging on muscle mass to body mass ratio and maximal running speed in rats. Body mass (a), muscle mass to body mass ratio (b) and maximal running speed (c) in the studied rat groups. Data is presented as mean ± SD. *p <0.05, **p <0.01 and *** p<0.001.

### Effects of intrinsic aerobic capacity on metabolic profiles of serum, muscle, and white adipose tissue

We used OPLS-DA to identify differences in the metabolic profiles between the HCRs and LCRs. At the age of 9 months (LCR-Y vs. HCR-Y), there were clear separations in serum (the goodness of the Q^2^ was 0.511 for OPLS-DA and the Q^2^ intercepts was –0.409 for the permutation test) and muscle (Q^2^ = 0.672, Q^2^ intercept = –0.321) metabolic profiles, but not in WAT (Q^2^ = −0.121, Q^2^ intercept = −0.072) (**Fig. S1, Table S1**). At the age of 21 months (LCR-O vs. HCR-O), there were clear separations in WAT (Q^2^ = 0.560, intercept = −0.513) and serum (Q^2^ = 0.258, intercept = −0.272) metabolic profiles, but not in muscle (Q^2^ = −0.182, intercept = −0.220) (**Fig. S2**, Horizontal axis describes the separation between groups and vertical axis describes the metabolic differences within the group).

Next, we compared the differences at the level of single metabolite between HCRs and LCRs with variable influence on projection analysis (VIP > 1.0 and p < 0.05 criteria **Fig. 2**). We identified several discriminating metabolites in serum and skeletal muscle of the young animals (LCR-Y vs. HCR-Y), but none in WAT (**Fig. 2**). In aged rats, LCRs had higher glutamate, isovalerylcarnitine and hexanoylcarnitine in serum, whereas no differences were found in muscle (**Fig. 2a-b**). In WAT, aspartate, arginine, sucrose, decanoylcarnitine, octanoylvcarnitine and kynurenate were downregulated in aged LCRs compared with HCRs (**Fig. 2c**). Only isovalerylcarnitine was upregulated in both young and aged LCRs compared with HCRs in serum (**Fig. 2a**).

**Figure. 2.**
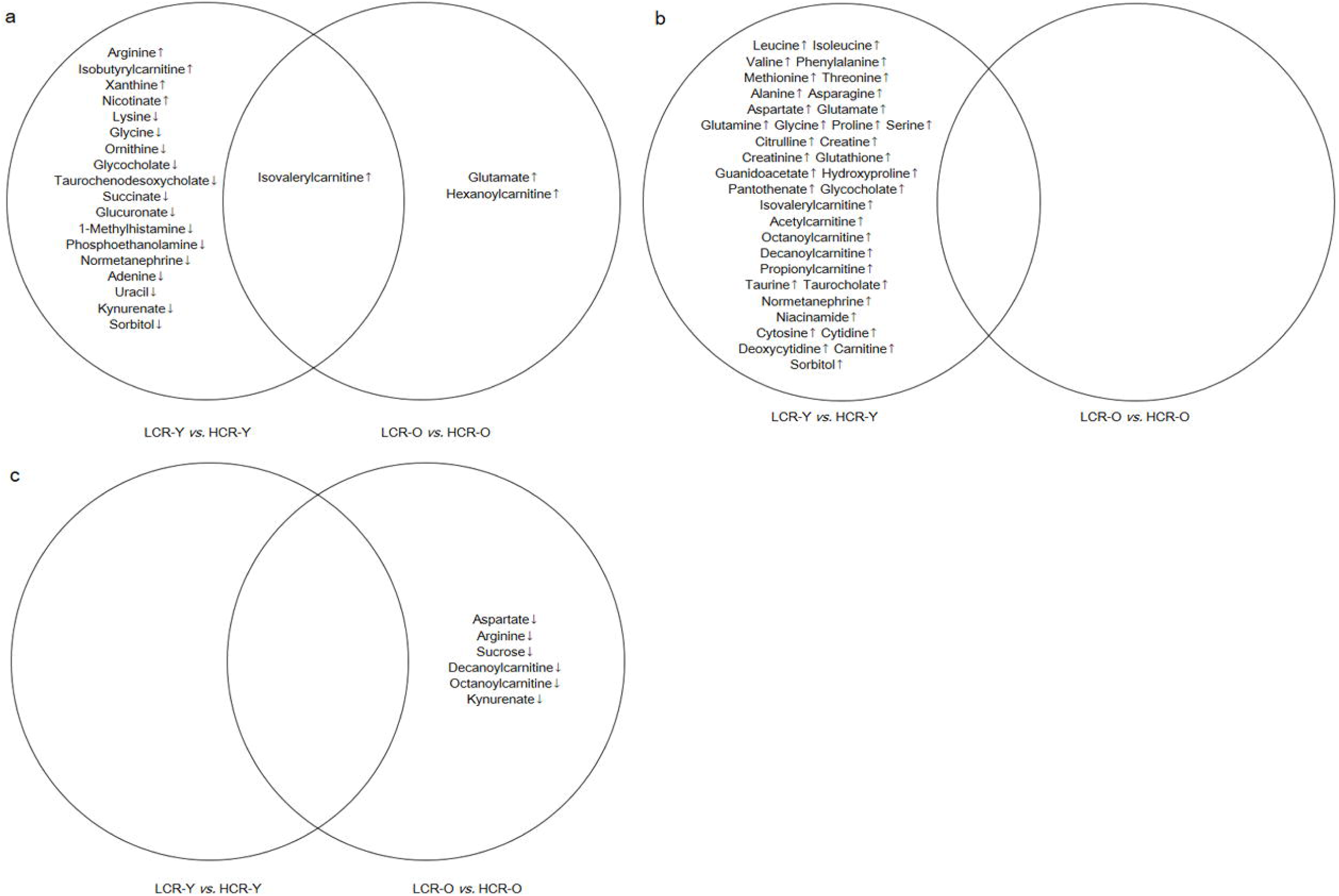
Venn diagrams of differentially expressed metabolites between LCR-Y *vs*. HCR-Y and LCR-O *vs*. HCR-O groups in serum (a), muscle (b) and WAT (c).

The key metabolic pathways that differed between the young LCRs and HCRs are illustrated in Figure 3. We found that in serum and muscle, amino acid metabolism differed most prominently between LCRs and HCRs at both age groups (**Table 1**). In addition, amino acid metabolism differed also in WAT in old LCRs and HCRs.

**Table 1.** Pathway analysis of LCR-Y vs. HCR-Y and LCR-O vs. HCR-O.

| Groups | Tissue | Pathway | FDR | Impact | Class |
| --- | --- | --- | --- | --- | --- |
| LCR-Y<br>vs.HCR-Y | Serum | Glycine, serine and threonine metabolism | 0.027 | 0.27 | Amino acid metabolism |
|  |  | Arginine and proline metabolism | 0.035 | 0.17 | Amino acid metabolism |
|  | Muscle | Alanine, aspartate and glutamate metabolism | 0.002 | 0.53 | Amino acid metabolism |
|  |  | Glycine, serine and threonine metabolism | 0.003 | 0.53 | Amino acid metabolism |
|  |  | D-Glutamine and D-glutamate metabolism | 0.002 | 0.50 | Metabolism of other amino acids |
|  |  | Taurine and hypotaurine metabolism | 0.004 | 0.43 | Metabolism of other amino acids |
|  |  | Glutathione metabolism | 0.002 | 0.36 | Metabolism of other amino acids |
|  |  | Arginine biosynthesis | 0.002 | 0.35 | Amino acid metabolism |
|  |  | Arginine and proline metabolism | 0.002 | 0.26 | Amino acid metabolism |
|  |  | Cysteine and methionine metabolism | 0.003 | 0.13 | Amino acid metabolism |
| LCR-O vs.<br>HCR-O | Serum | D-Glutamine and D-glutamate metabolism | 0.027 | 0.50 | Metabolism of other amino acids |
|  |  | Alanine, aspartate and glutamate metabolism | 0.027 | 0.20 | Amino acid metabolism |
|  |  | Arginine biosynthesis | 0.027 | 0.12 | Amino acid metabolism |
|  | WAT | Alanine, aspartate and glutamate metabolism | 0.033 | 0.22 | Amino acid metabolism |

**Figure. 3.**
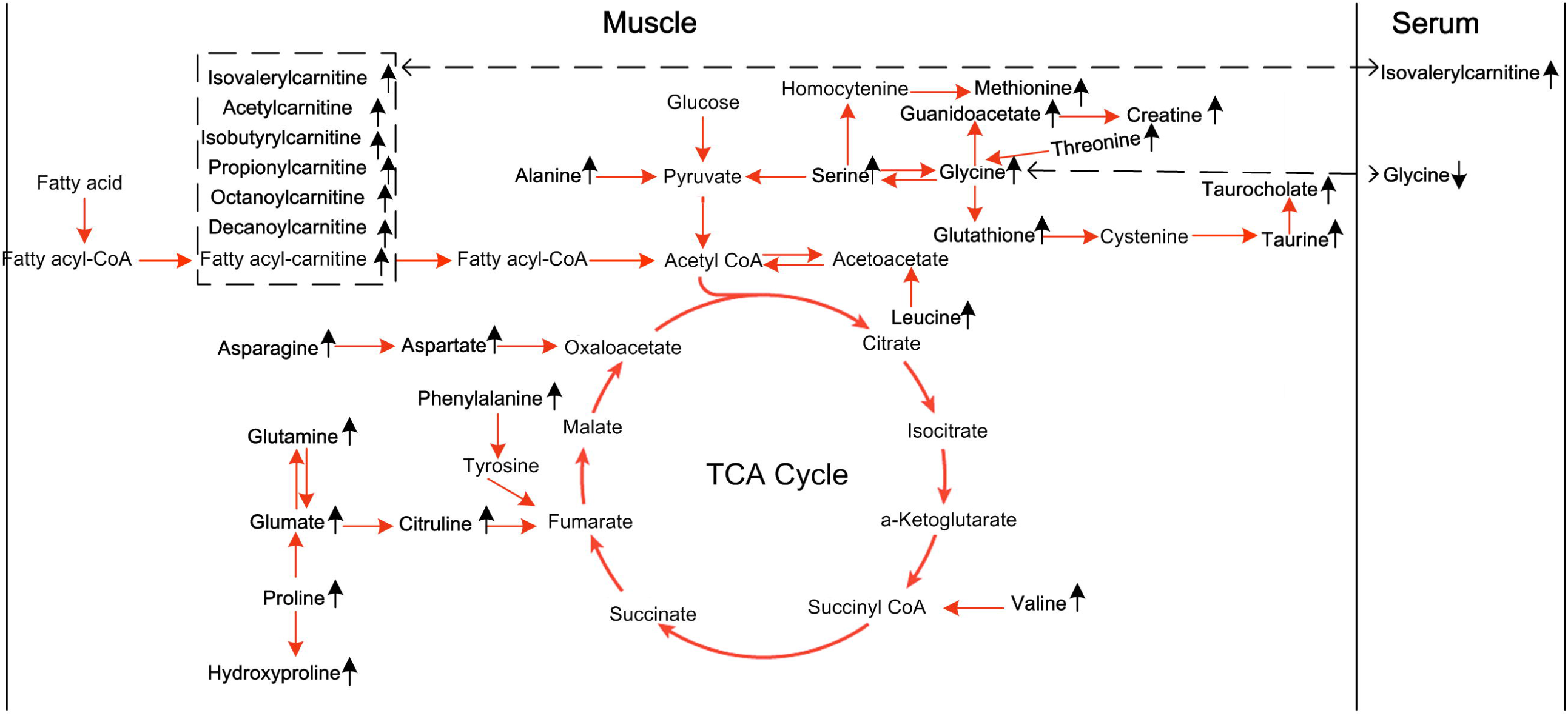
A schematic overview of the key metabolic pathways and the biochemical linkages among metabolites detected in the LCR-Y compared with HCR-Y. The significantly altered metabolites are bolded. The black arrows represent up/down-regulated metabolites in the LCR-Y compared with the HCR-Y. The red arrows represent metabolic pathway link. The dotted line represents the links of metabolites between muscle and serum.

### Effects of aging on metabolic profiles of serum, muscle, and white adipose tissue

To better understand the effect of aging on metabolic profiles of LCRs and HCRs, we compared the metabolites of serum, skeletal muscle and WAT between the young (9 months old) and old (21 months old) rats (**Fig. 4**). The metabolic profiles of all the studied tissues were clearly separated between the young and old in both rat lines (**Fig. S3, Fig. S4 and Table S3**). There were several common metabolites that changed with aging in both LCRs and HCRs in serum, muscle, and WAT (**Fig. 4**). However, we observed also distinct aging patterns in metabolites between LCRs and HCRs. In serum, lysine, 2-aminoisobutyrate, ADMA, pantothenate, normetanephrine and niacinamide were downregulated in HCRs whereas GABA, taurochenodeoxycholate, succinate, taurine, adenosine and choline were upregulated in LCRs (**Fig. 4a**). In muscle of HCRs, glutathione, glutamate, GABA, decanoylcarnitine, normetanephrine, hypoxanthine and AMP were upregulated whereas several amino acids (e.g. leucine, valine, isoleucine, phenylalanine, tyrosine) were downregulated with aging in LCRs. In WAT, niacinamide, citrulline and glutathione were upregulated in LCRs, whereas numerous metabolites were upregulated in aged HCRs (**Fig. 4c**). The predominant pathways that differed between young and old HCRs and LCRs in all three tissues were amino acid metabolism, lipid metabolism, nucleotide metabolism, and cofactors and vitamins metabolism (**Fig. 5, Table 2**).

**Table 2.**
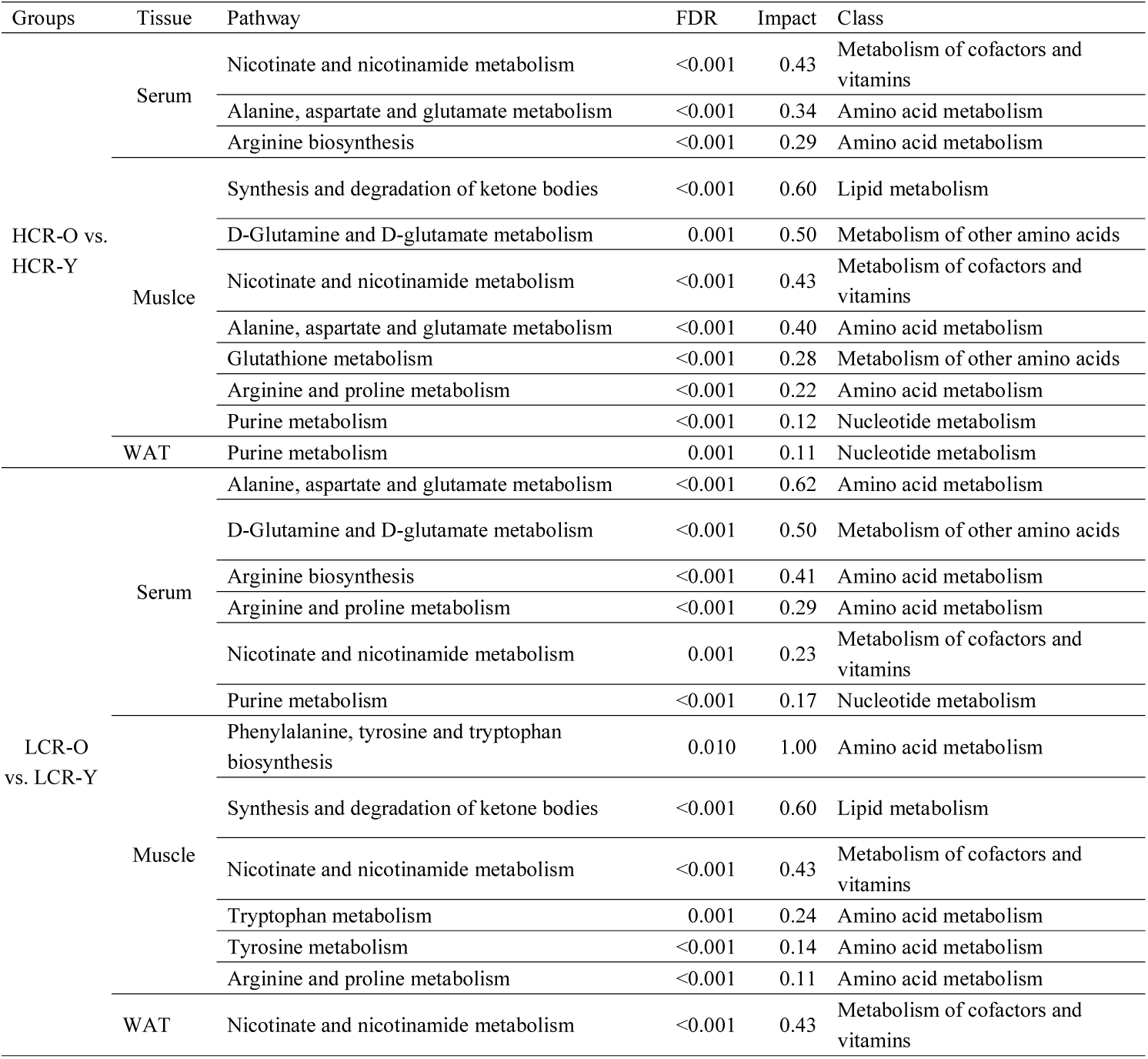
Pathway analysis of HCR-O vs. HCR-Y and in LCR-O vs. LCR-Y.

**Figure. 4.**
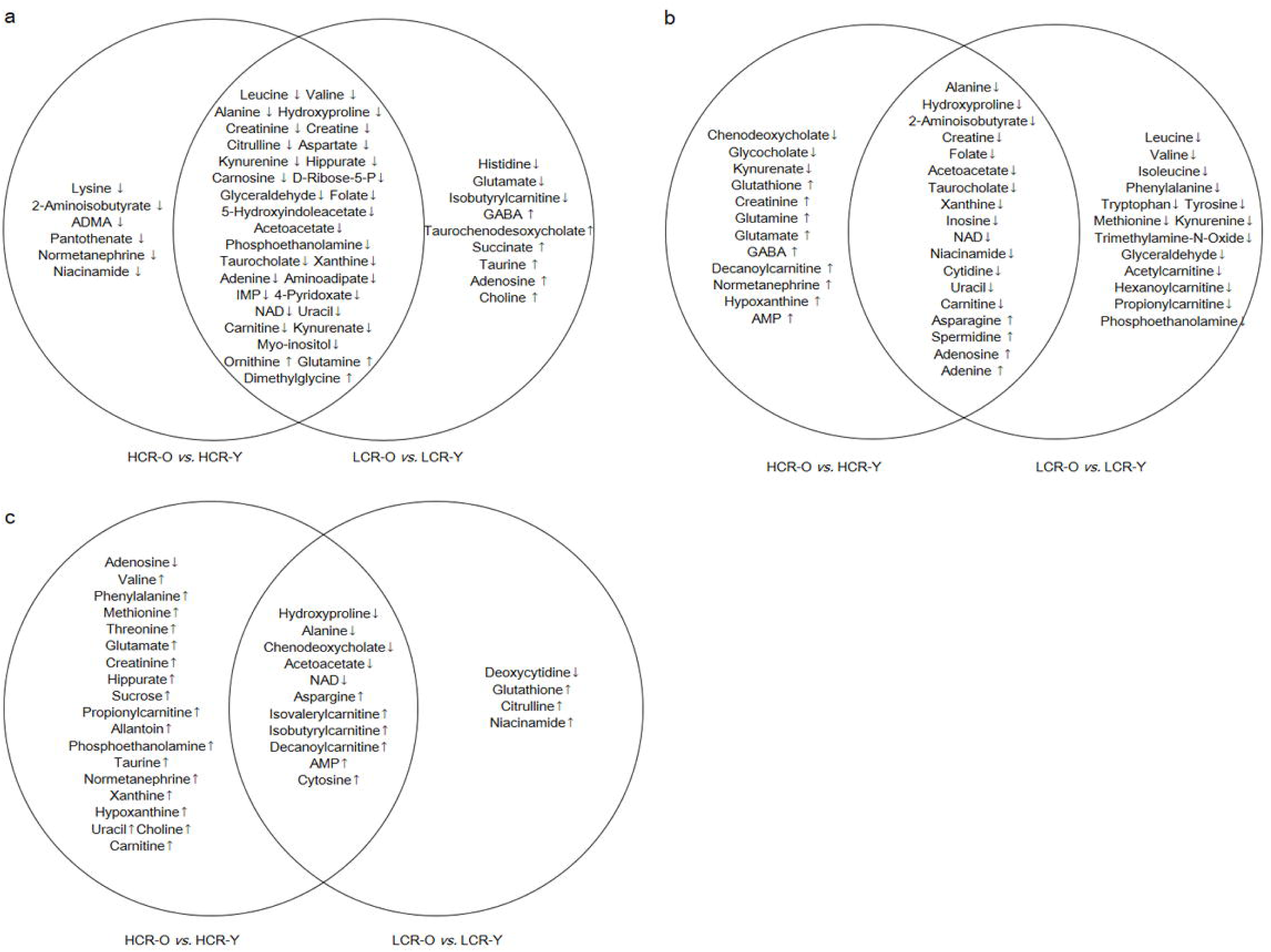
Venn diagrams of significantly different metabolites between HCR-O *vs*. HCR-Y groups and LCR-O *vs*. LCR-Y groups in serum (a), muscle (b) and WAT (c).

**Figure. 5.**
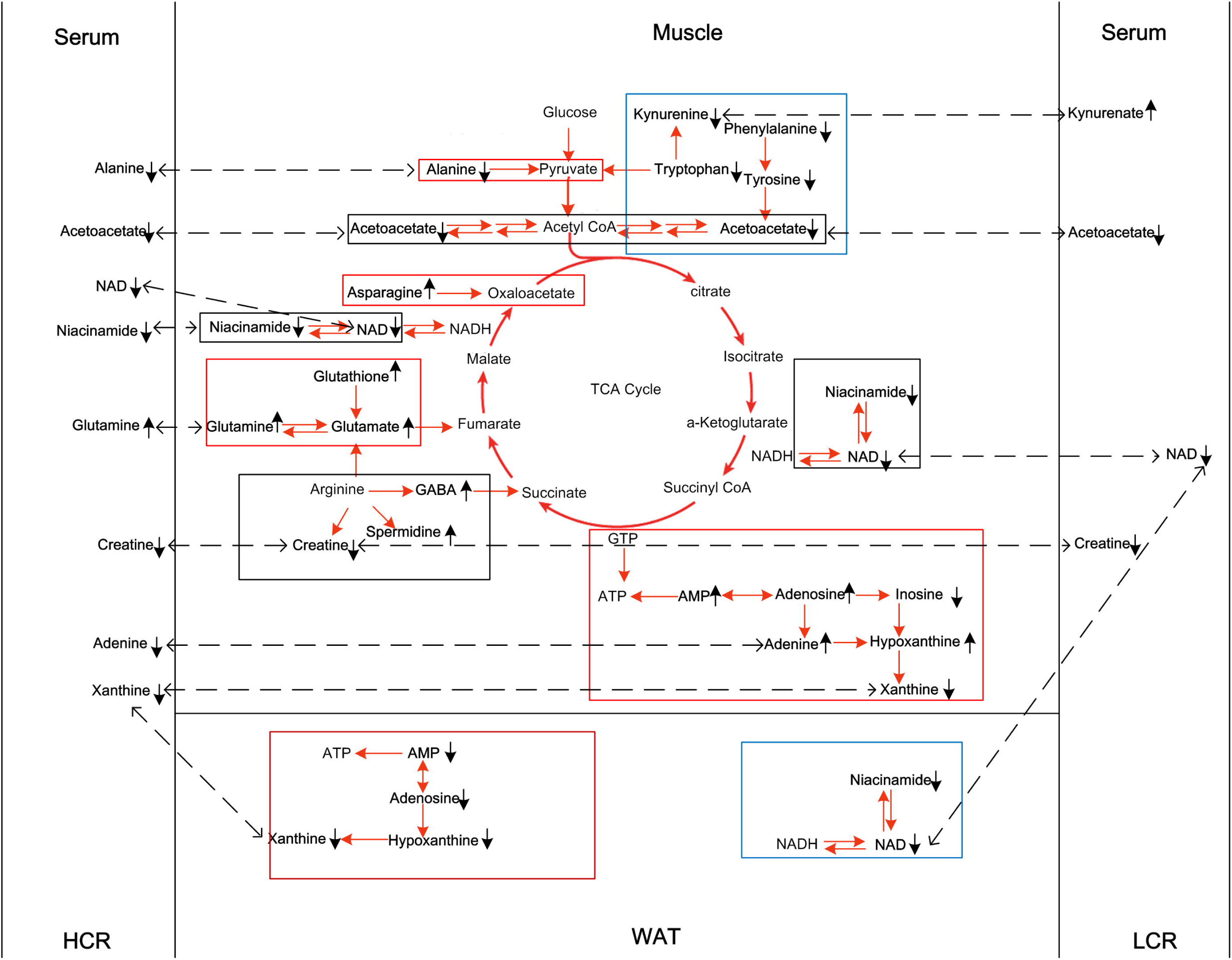
A schematic overview of the most predominantly differential metabolic pathways and the biochemical linkages among metabolites detected in old compared with young rats. The significantly altered metabolites are bolded. The black arrows represent up/down-regulated metabolites in the old rats compared with the young rats. The red arrows represent metabolic pathway links. The dotted line represents the links of metabolites between muscle and serum, and between WAT and serum. Pathways in black box represent the altered pathways in both LCRs and HCRs during aging; pathways in red box represent specific altered pathways in HCR, and pathways in blue box were the specific altered pathways in LCR.

### The interaction of aerobic capacity and aging on metabolites

The interaction between aerobic capacity and aging on metabolites was assessed using regression models. We found that high aerobic capacity and aging interactively affected alanine and carnosine in serum (p < 0.05, **Table 3**), acetoacetate, isovalerylcarnitine,and niacinamide in muscle (**Table 3**) and adenosine and chenodeoxycholate in WAT (**Table 3**). These changes were driven by HCRs.

**Table 3.** The unstandardized regression coefficient of metabolites which were interactively affected by aerobic capacity and aging in serum, muscle and WAT.

| Tissue | Metabolites | Aerobic capacity | Age |  |
| --- | --- | --- | --- | --- |
|  |  |  | Young | Old |
| Serum | Alanine | Low | 0.05 | 0.01 |
|  |  | High | 0.24 | -0.36 |
|  | Carnosine | Low | 4.50 | 12.15 |
|  |  | High | 59.13 | -313.57 |
| Muscle | Acetoacetate | Low | 0.41 | 0.82 |
|  |  | High | -0.05 | -5.60 |
|  | Isovalerylcarnitine | Low | 243.77 | -61.41 |
|  |  | High | 1553.85 | -3498.57 |
|  | Niacinamide | Low | -0.64 | 0.40 |
|  |  | High | 0.11 | -3.67 |
| WAT | Adenosine | Low | 0.03 | 0.13 |
|  |  | High | -0.74 | 142.65 |
|  | Chenodeoxycholate | Low | 0.03 | 3.75 |
|  |  | High | 0.06 | 31.02 |

## DISCUSSION

The present study showed that aerobic capacity and aging have different impacts on the metabolic profiles of serum, skeletal muscle, and white adipose tissue (WAT). According to our hypothesis, at a young age, metabolites linked to amino acid metabolism differed with aerobic capacity in serum and muscle, whereas no differences were observed in WAT. Interestingly, in aged animals, most prominent changes in metabolites occurred in WAT of the HCRs. Our study highlights that aerobic capacity is predominantly associated with more efficient amino acid metabolism in muscle and emphasizes the importance of WAT during aging. Noticeably, the interactive effects of aging and aerobic capacity on energy metabolism-related metabolites were largely driven within HCRs, reflecting the importance of inherited aerobic capacity in the processes of aging.

### High aerobic capacity is associated with more efficient amino acid metabolism in muscle

Skeletal muscle is an important tissue contributing both to aerobic capacity and whole-body metabolism [29]. According to our hypothesis, several metabolites linked to amino acid metabolism were downregulated in serum and upregulated in skeletal muscle of the young LCRs compared with HCRs (**Fig. 2**), indicating that HCRs have more efficient amino acid metabolism in skeletal muscle. Our observation is in line with previous findings, where young (3-4.5 months of age) HCRs were found to oxidize branched-chain amino acids more efficiently compared to LCRs of the same age [17].

In our study, a total of 11 amino acids (isoleucine, valine, phenylalanine, methionine, threonine, alanine, asparagine, aspartate, glutamate, glutamine, and glycine) were higher in muscle in young LCRs compared to HCRs (F**ig. 2b**). Previous study by Overmyer *et al*. found that muscle BCAAs (leucine, isoleucine, and valine) were lower in HCRs than LCRs after 10 min of speed-ramped treadmill running [17]. These changes were paralleled by a fall in plasma BCAAs in HCRs, indicating increased utilization of BCAAs during exercise. In our study, the serum level of arginine was higher, but levels of lysine, glycine and ornithine were lower in serum of young LCRs compared with HCRs (**Fig. 2a**). It seems that during resting at fasted state, HCRs do not utilize the amino acids from serum, even though the metabolites linked to amino acid metabolism in muscle are low. In addition, several metabolites linked to BCAA and fatty-acid metabolism (i.e. isovalerylcarnitine, acetylcarnitine, propionylcarnitine, octanoylcarnitine and decanoylcarnitine) were higher in muscle in young LCRs compared with HCRs (**Fig. 2a**), indicating more efficient fatty-acid oxidation in skeletal muscle of HCRs, as shown before [17, 24]. The low level of the glucogenic amino acids accompanied by lower level of several fatty-acid metabolites indicate higher tricarboxylic acid (TCA) cycle turnover in HCRs (**Fig. 3**), supporting previous findings [17, 18, 24]. Glucogenic amino acids can serve as anaplerotic substrates, which can directly feed into the TCA cycle [30]. The higher TCA cycle in HCRs was responsible for the higher resting energy expenditure [31], higher body temperature [18], and better exercise performance [32] c ompared with LCRs.

On the other hand, higher levels of metabolites linked to BCAA and fatty-acid metabolism in LCRs suggest impaired mitochondrial energy production in LCRs [17]. Elevated levels of acyl carnitines have been observed also in a model of type 2 diabetes fatty Zucker rats that have significantly decreased activity of BCAA catabolism [33]. Furthermore, the accumulation of incompletely oxidized lipids contribute to skeletal muscle insulin resistance [34]. These results further support the previous finding that the high risk of metabolic disorders in LCRs is associated with impaired BCAA metabolism [17]. Hence increasing aerobic capacity to restore the BCAA and β-oxidation in the muscle could reduce insulin resistance. Recent studies have shown that skeletal muscle mitochondrial dysfunction induced by oxidative stress led to an increase in biosynthetic serine/1-carbon/glycine pathway and an elevation in glycine, serine, and glutathione in skeletal muscle [35, 36]. The high levels of glycine, serine and glutathione indicate mitochondrial dysfunction in the LCRs, which may reflect a compensatory activation of stress signaling network of glycine/serine metabolism that preserves cellular function induced by increased mitochondrial stress.

Interestingly, we found that the level of glycine was low in the serum but high in the muscle in LCRs (**Fig. 2**). Glycine can act as an extracellular carrier of various endogenous and xenobiotic metabolites, including derivatives of BCAAs and ß-oxidation intermediates [37]. The elevated level of BCAAs and fatty acyl carnitines and increased mitochondrial stress in the muscle of LCRs may be considered to be responsible for lowering glycine levels in the serum. Here, we suggest that the low level of glycine in the serum may reflect the increased need of this amino acid in the skeletal muscle metabolism.

Compared with other metabolically active tissues, the oxidative capacity of WAT is relatively low. For instance, O_2_ consumption per kg wet weight in WAT is about one-tenth of that of the resting skeletal muscle [38]. We found no significant differences in metabolite concentrations of WAT between young LCRs and HCRs (**Fig. 2c**). This agrees well with the previous studies showing that intrinsic aerobic capacity does not influence the mitochondrial protein abundance and expression or metabolic activity in WAT of the young rats [22, 39].

When examining the old animals, only three metabolites (glutamate, isovalerylcarnitine and hexanoylcarnitine) were upregulated in serum of LCRs compared with HCRs. Increased concentrations of isovalerylcarnitine and hexanoylcarnitine have been previously reported in type 2 diabetes [40-42], which is in line with previous results showing increased blood glucose and insulin in LCRs compared with HCRs [17, 24]. Interestingly, there were six metabolites downregulated in WAT in old LCRs compared with HCRs (F**ig. 2c**). Three of the metabolites were linked to amino acid metabolism (aspartate, arginine and kynurenate) and two to fatty-acid metabolism (decanoylcarnitine and octanoylcarnitine). Among these three metabolites, kynurenate increases energy utilization by stimulating thermogenic gene expression in adipose tissue [43]. The elevated kynurenate in HCRs supports our previous study where HCRs had higher body temperature [18] than LCRs. However, the downregulation of two fatty acyls indicate higher fatty acid metabolism in WAT in old LCRs compared with HCRs whereas no differences were observed in skeletal muscle (**Fig. 2b**).

### Aerobic capacity affects the metabolic profile with aging in serum, muscle and especially WAT

Our results revealed several common altered metabolites in serum, muscle, and WAT (**Fig. 4**). However, we also observed distinct aging patterns in both examined tissues as well as in serum. In HCRs, the metabolites that differed from LCRs were downregulated in old compared with the young rats and linked mainly to amino acid and vitamin metabolism in the serum, (**Fig 4a and Table 2**). In contrast, most of the metabolites altered by aging solely in LCRs were upregulated in old animals, yet were similarly linked mainly to amino acid, vitamin and nucleotide metabolism (**Fig 2a, Table 2**). Amino acid availability is critical in the regulation of muscle protein metabolism [44]. With aging, the serum concentrations of amino acids decrease. This is mainly related to the reduction of dietary protein intake and alterations in gluconeogenesis and the function of urea cycle [45, 46]. A previous study examined the metabolomics profile of plasma associated with aging in HCRs/LCRs [16]. Metabolomic profiles of young (13 month) and old (26 month) male HCR and LCR rats were analysed using proton nuclear magnetic resonance spectroscopy. A strong profile separation occurred in old and LCRs, whereas young and HCRs were found less predictive [16]. The reason for the different observations may be due to differences in sex, generation (Falavigna *et al*. used generations 17 and 19), fasting time and euthanasia sacrifices method. Here we showed the decreased levels of serum amino acids with aging that may reduce the basal rate of muscle protein synthesis, thereby contributing to loss of muscle mass (**Fig 1b and Fig 4**a). Our findings further indicate that at old age, HCRs are able to more efficiently utilize metabolites from serum, whereas in LCRs the opposite is true.

In muscle, aging leads to upregulation of several metabolites in HCRs and downregulation of several metabolites in LCRs that were mainly linked to amino acid and lipid metabolism (**Fig 4b, Table 2**). Our results indicate that certain metabolic pathways activate in HCRs with aging possibly combating the unfavorable changes in skeletal muscle metabolism and loss of muscle mass that occurs with aging [47]. In LCRs, many amino acids (leucine, isoleucine, valine, phenylalanine, tryptophan, tyrosine, and methionine) as well as fatty-acid metabolites (acetylcarnitine, hexanoylcarnitine and propionylcarnitine) were downregulated in old rats (**Fig. 4b**), suggesting a decreased TCA turnover with aging.

In WAT several metabolites were upregulated in old HCRs while only a few were upregulated in LCRs (**Fig. 4c**). In WAT of old rats, decreased level of acetoacetate but increased levels of isovalerylcarnitine, isobutyrylcarnitine and decanoylcarnitine suggest dysfunctional mitochondrial β-oxidation [34]. This observation is in agreement with previous studies showing that adipose tissue plays a central role in the development of insulin resistance, metabolic dysfunction, inflammation, and impaired regenerative capacity during aging [48-50]. Intriguingly, aging seemed to more prominently affect the metabolite levels of WAT in HCRs, where several amino acids (e.g. valine, phenylalanine, methionine and threonine) were upregulated (**Fig 4c**), indicating a decrease in amino acid utilization in WAT tissue with aging.

### Aerobic capacity and aging interactively affect metabolites in serum, muscle, and WAT

Previous human studies have shown that high aerobic capacity reduces symptoms of age-related disorders, including obesity, diabetes, inflammation and cardiovascular diseases [51, 52], suggesting that aerobic capacity and aging have interactive effects on metabolism. We found seven metabolites were interactively affected by aerobic capacity and aging (**Table 3**). The observed interaction between high aerobic capacity and aging is reasonable due to the significant decline in aerobic capacity with aging specifically in HCRs (**Fig. 1c**). No age effect on metabolites and aerobic capacity in LCRs was found, which is likely due to the low initial aerobic capacity, which did not significantly decline with age.

Aging and high aerobic capacity interactively decreased the levels of alanine and carnosine in serum (**Table 3**). A previous study has shown that β-alanine supplementation increased exercise capacity in middle aged individuals [53]. Alanine is also a crucial intermediate of muscle energy metabolism and regulates inter-organ glucose homeostasis via the glucose-alanine cycle [54]. Alanine activates the AMPK pathway and acutely reduces blood glucose to restore energy homeostasis [55]. In addition, carnosine is synthesized from alanine and is able to improve myofibril contractility via enhancing sarcomere sensitivity to calcium, as well as to maintain pH homeostasis in muscle, possibly contributing to endurance capacity [56, 57]. Efficient energy production is critical for the maintenance of high aerobic capacity. The decreased concentration of energy metabolism-related metabolites with aging (acetoacetate, isovalerylcarnitine, and niacinamide) in the skeletal muscle of HCRs may reflect the importance of inherited aerobic capacity in the processes of aging. Niacinamide is a precursor of a key coenzyme, nicotinamide adenine dinucleotide (NAD), which coordinates several reactions related to energy metabolism, correct folding of proteins as well as DNA repair and mitochondrial maintenance [58, 59]. Moreover, NAD has emerged as a potential target that counteracts metabolic disturbances and the mitochondrial dysfunction that is mediated through sirtuins [60]. In agreement with this finding we have shown that HCRs have higher sirtuin 3 level in skeletal muscle [23].

In WAT, decreased adenosine with aging in HCRs was observed. A previous study showed that adenosine is a potent inhibitor of lipolysis via inhibiting cAMP accumulation [61]. Adenosine is also an insulin sensitizer and therefore stimulates glucose transport and oxidation [62] which indicate that high aerobic capacity may preserve the insulin sensitivity decline with aging. Furthermore, chenodeoxycholate was affected by both aging and aerobic capacity (**Table 3**). Decreased chenodeoxycholate is attributed to the activation of critical inflammatory regulators such as NF-kB, and ameliorate insulin resistance by modulation of adipokines [63]. The age related adipose tissue dysfunction including peripheral insulin resistance and inflammation may be associated with adenosine and chenodeoxycholate. Hence, enhancing the aerobic capacity by exercise in order to recover these metabolites may decrease the risk of age-related metabolic disorders.

### Conclusion

Our results suggest that intrinsic aerobic capacity drives differences in metabolism mainly in muscle, not in WAT. Our results support the notion that impaired BCAA and fatty acid metabolism in the muscle may relate to the high risk of metabolic disorders and early mortality previously shown in LCRs. The interactive effects of aging and aerobic capacity on energy metabolism-related metabolites were driven by HCRs, reflecting the importance of inherited aerobic capacity in the processes of aging. Furthermore, dysfunctional mitochondrial β-oxidation in WAT may be one of the mechanisms related to aging. Future studies are needed to determine whether improved aerobic capacity by training could restore the observed aging related metabolic changes.

---

**Supplementary figure 1**. Score plot of OPLS-DA in serum (a), muscle (c) and WAT (e) from LCR-Y and HCR-Y groups, and a 200 times permutation test of OPLS-DA models for serum (b), muscle (d) and WAT (f). The blue circle represents HCR-Y group, and yellow circle represents LCR-Y group.

**Supplementary figure 2**. Score plot of OPLS-DA in serum (a), muscle (c) and WAT (e) from LCR-O and HCR-O groups, and a 200 times permutation test of OPLS-DA models for serum (b), muscle (d) and WAT (f). The green circle represents HCR-O group, and red circle represents LCR-O group.

**Supplementary figure 3**. Score plot of OPLS-DA in serum (a), muscle (c) and WAT (e) from HCR-O and HCR-Y groups, and a 200 times permutation test of OPLS-DA models for serum (b), muscle (d) and WAT (f). The blue circle represents LCR-Y group, and green circle represents LCR-O group.

**Supplementary figure 4**. Score plot of OPLS-DA in serum (a), muscle (c) and WAT (e) from LCR-O and LCR-Y groups, and a 200 times permutation test of OPLS-DA models for serum (b), muscle (d) and WAT (f). The yellow circle represents LCR-Y group, and red represents LCR-O group.

**Supplementary Table 1.**
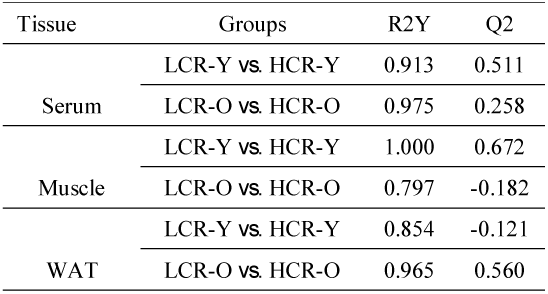
The R^2^Y and Q^2^ of OPLS-DA model for aerobic capacity effect.

| Tissue | Groups | R2Y | Q2 |
| --- | --- | --- | --- |
| Serum | LCR-Y vs. HCR-Y | 0.913 | 0.511 |
|  | LCR-O vs. HCR-O | 0.975 | 0.258 |
| Muscle | LCR-Y vs. HCR-Y | 1.000 | 0.672 |
|  | LCR-O vs. HCR-O | 0.797 | -0.182 |
| WAT | LCR-Y vs. HCR-Y | 0.854 | -0.121 |
|  | LCR-O vs. HCR-O | 0.965 | 0.560 |

**Supplementary Table 2.**
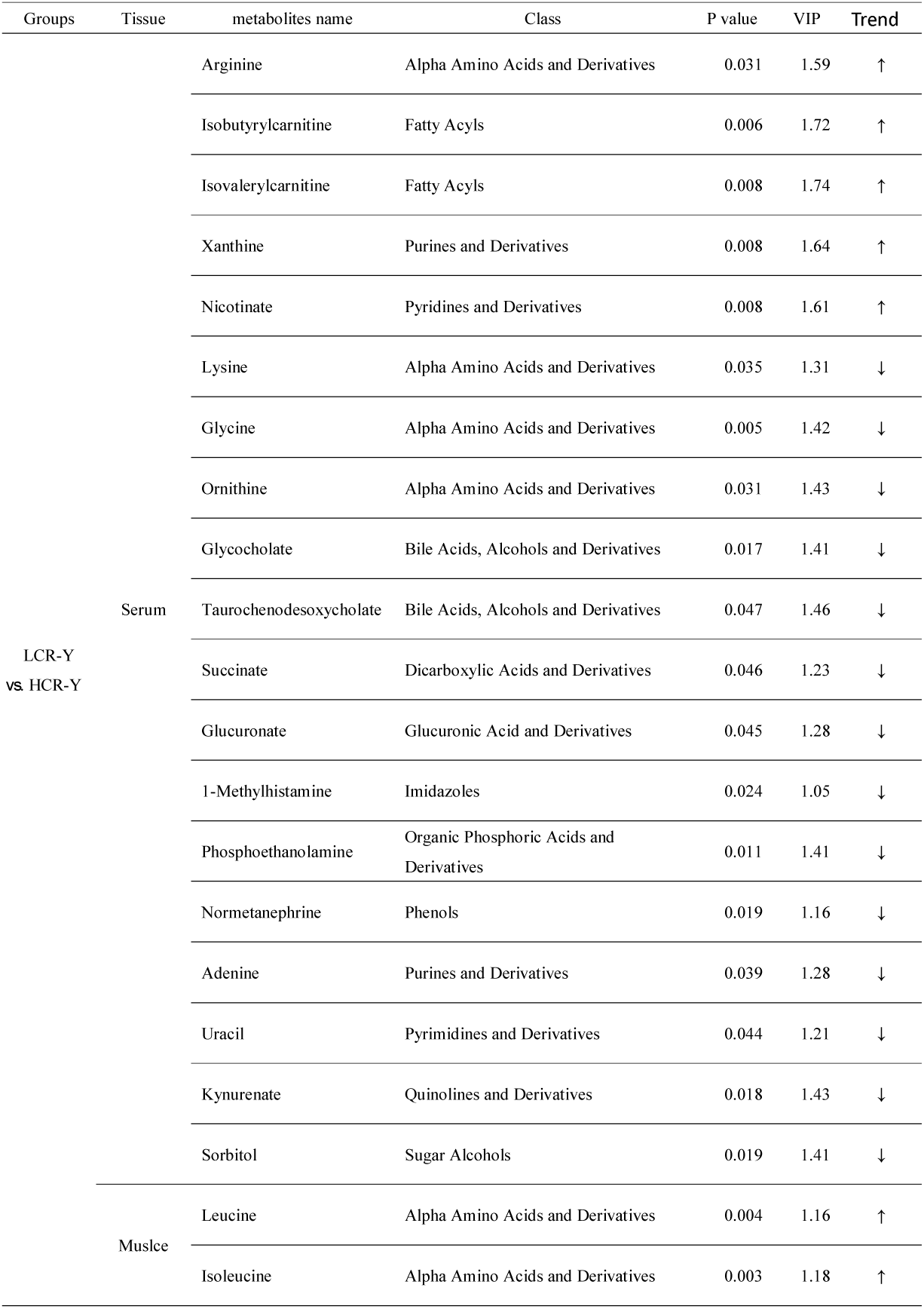
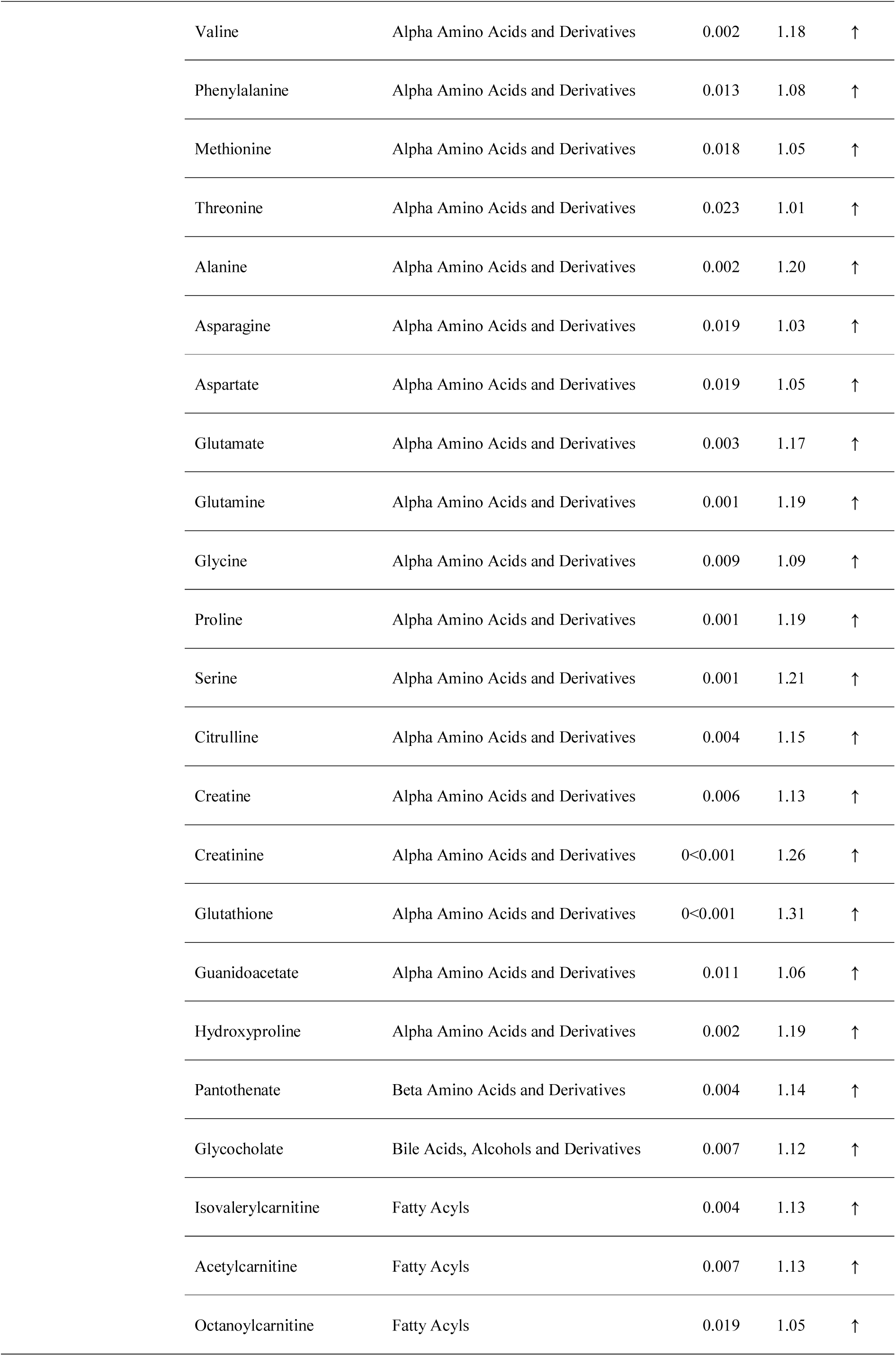
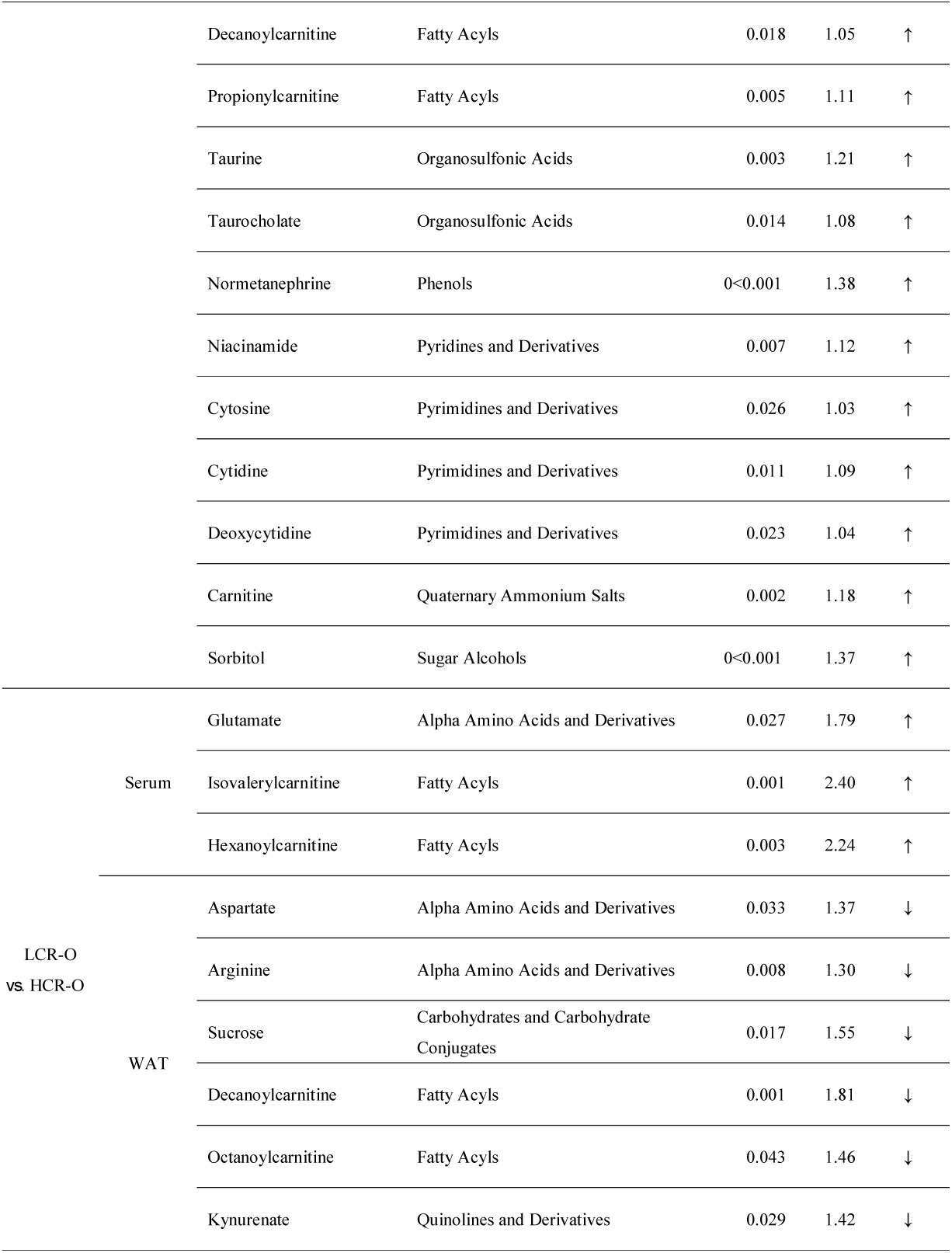
Significantly different metabolites induced by aerobic capacity.

**Supplementary Table 3.** The R^2^Y and Q^2^ of OPLS-DA model for aging effect.

| Tissue | Groups | R2Y | Q2 |
| --- | --- | --- | --- |
| Serum | HCR-O vs. HCR-Y | 1.000 | 0.974 |
|  | LCR-O vs. LCR-Y | 1.000 | 0.943 |
| Muscle | HCR-O vs. HCR-Y | 0.995 | 0.972 |
|  | LCR-O vs. LCR-Y | 0.975 | 0.959 |
| WAT | HCR-O vs. HCR-Y | 0.999 | 0.975 |
|  | LCR-O vs. LCR-Y | 0.998 | 0.946 |

**Supplementary Table 4.**
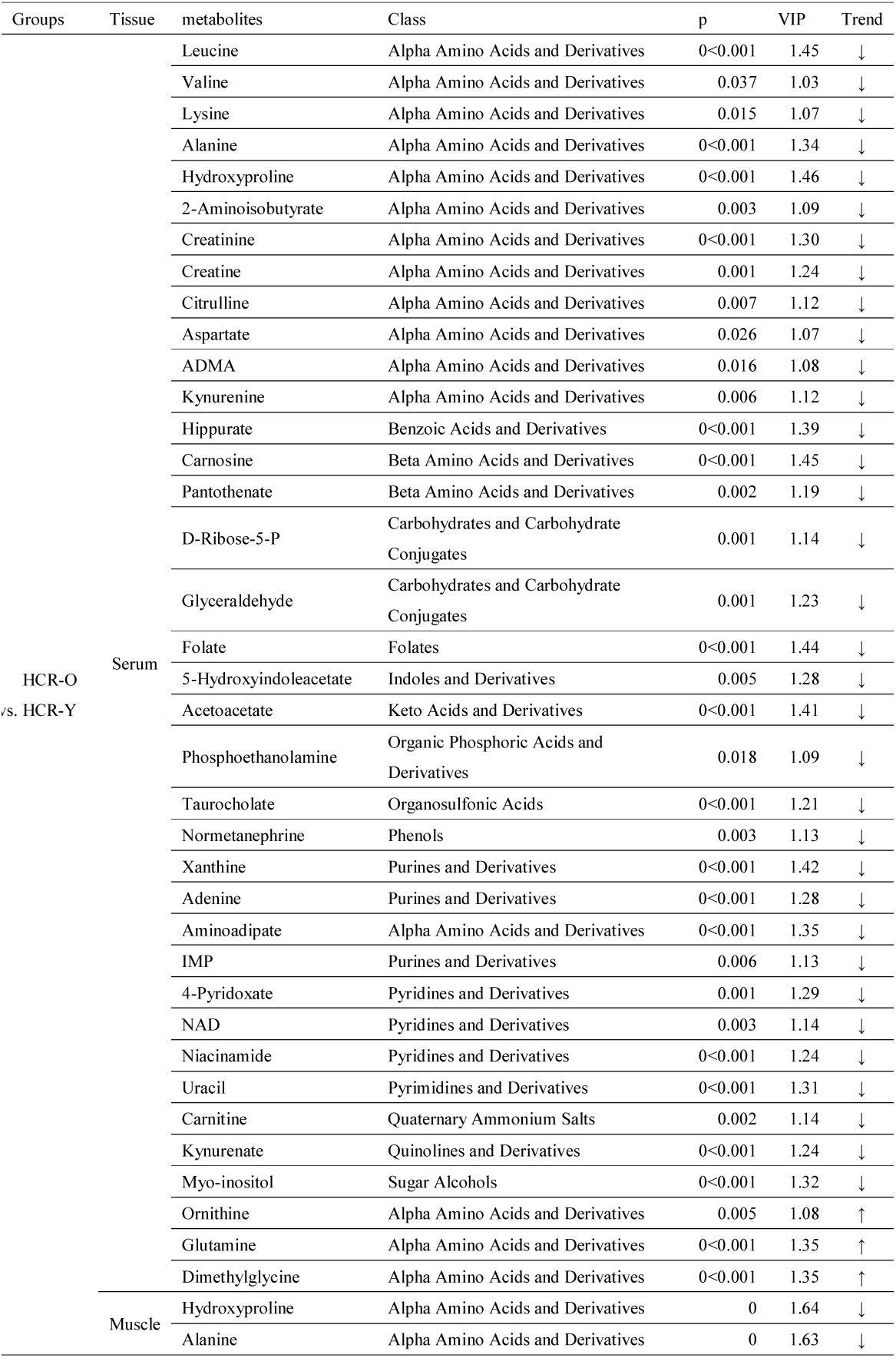
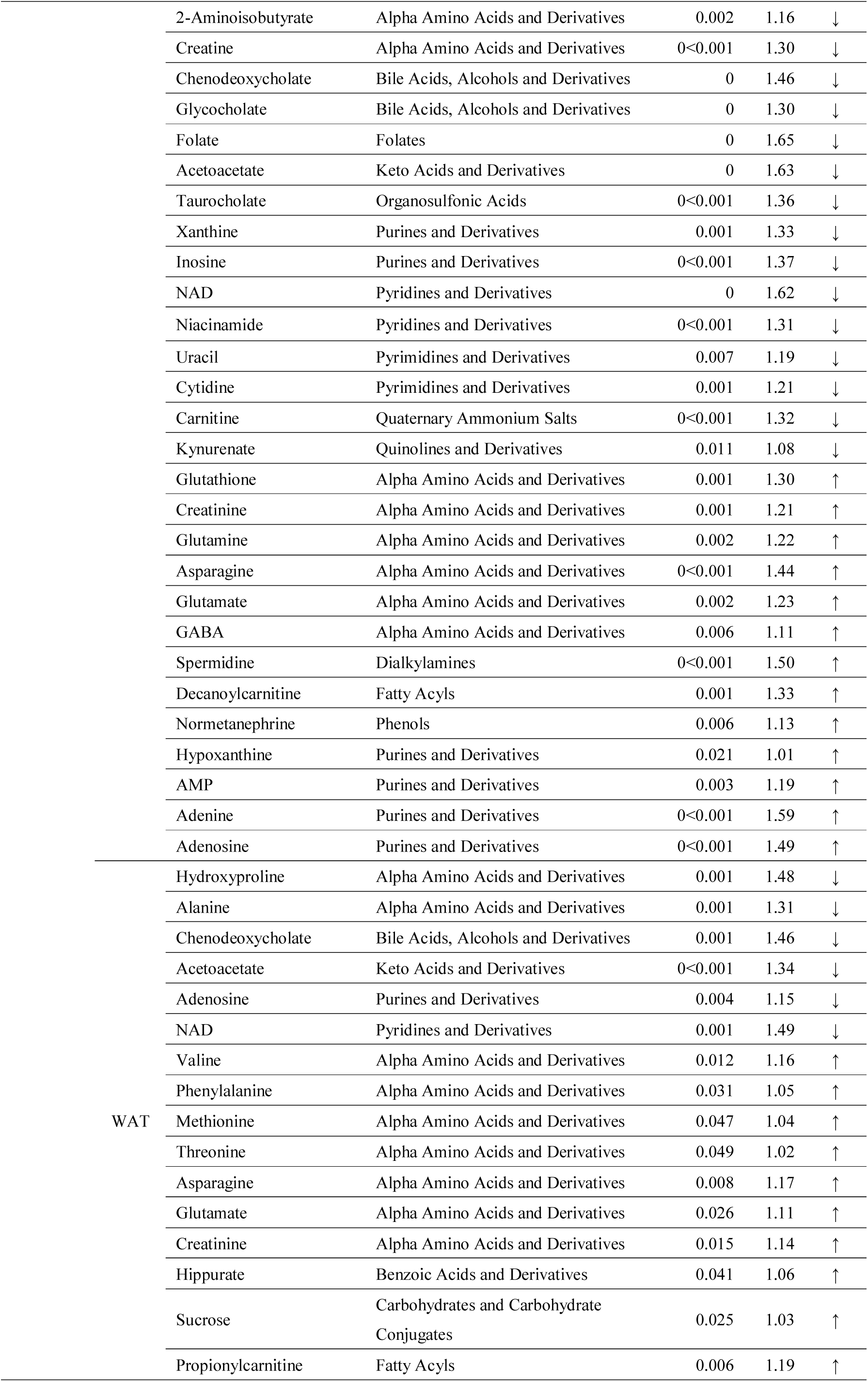
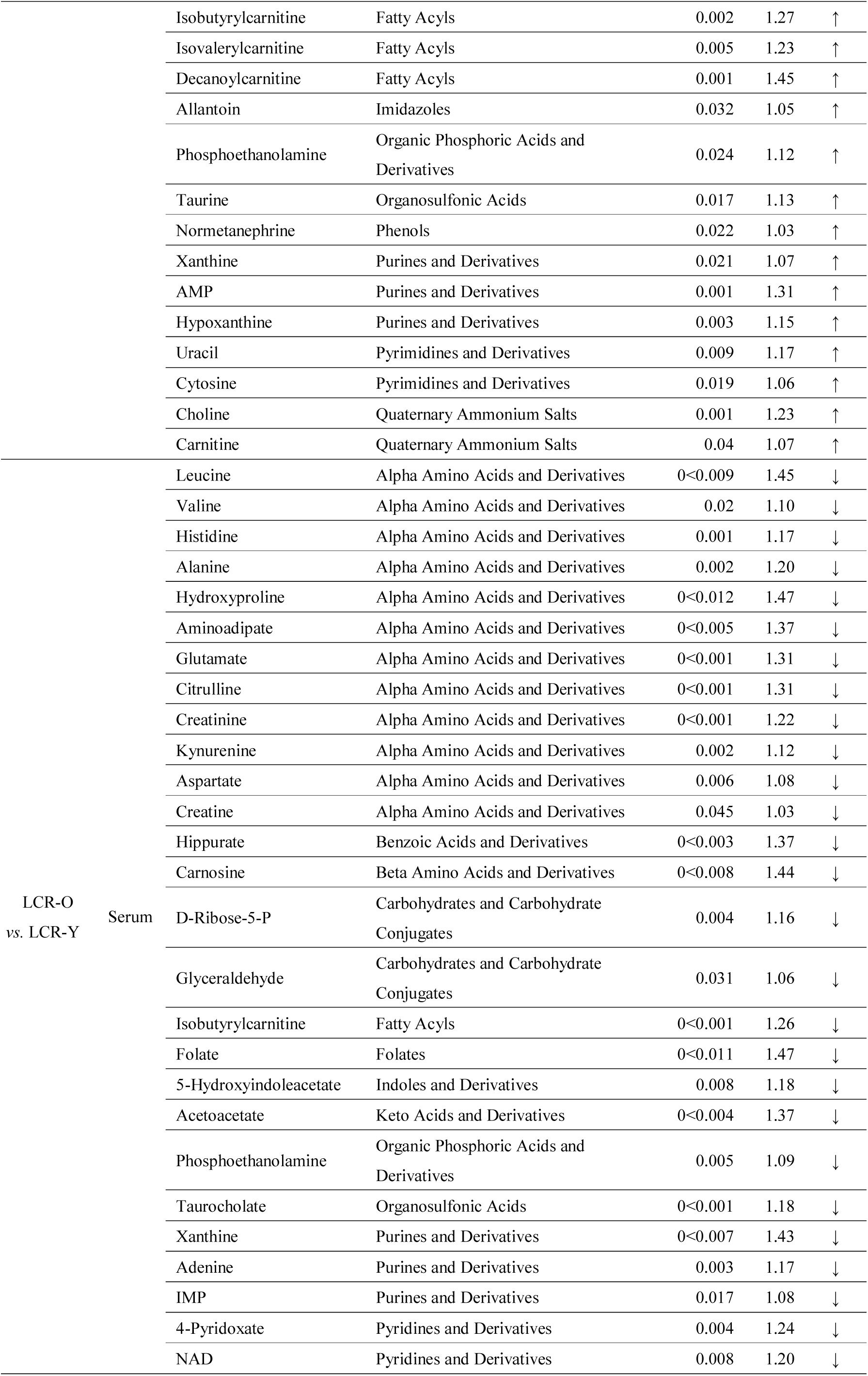
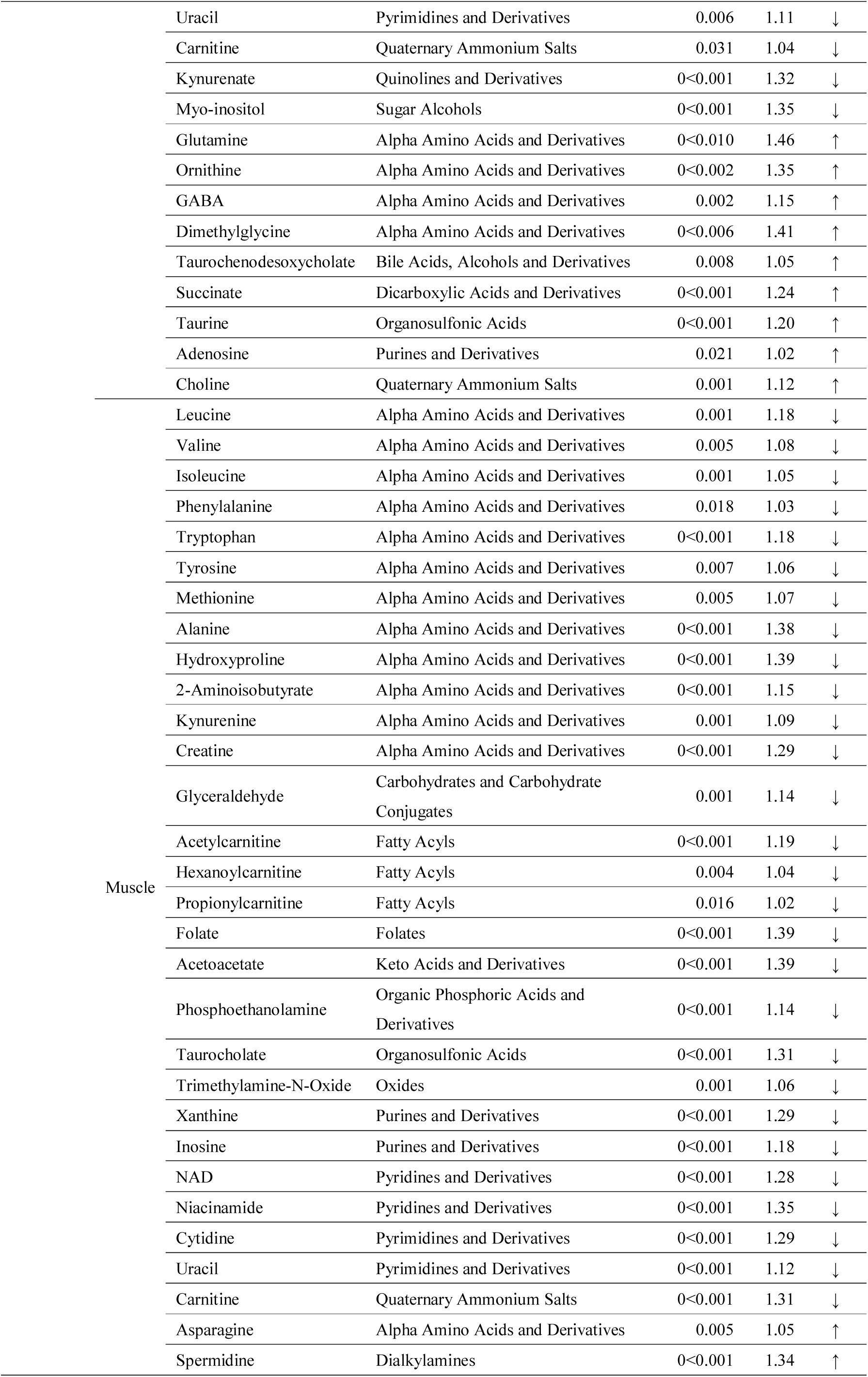
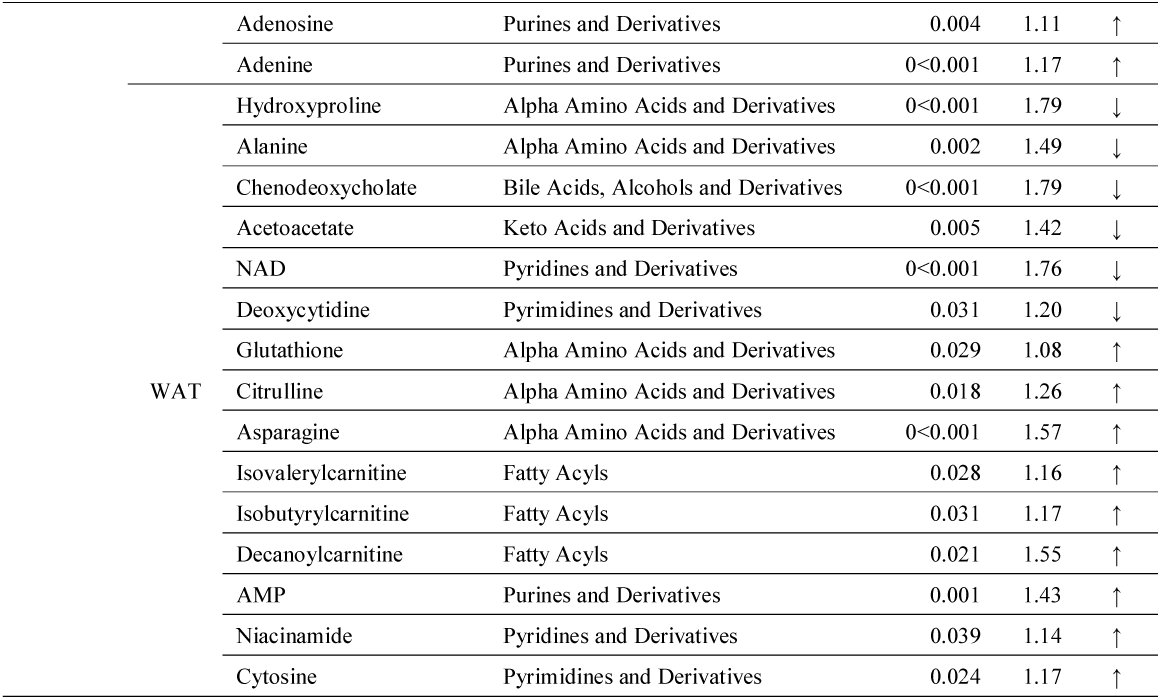
Significantly different metabolites induced by aging

