## Supplementary figures and images for "Interactive effects of aging and aerobic capacity on energy metabolism-related metabolites of serum, skeletal muscle, and white adipose tissue"

### Supplemental Figure 1

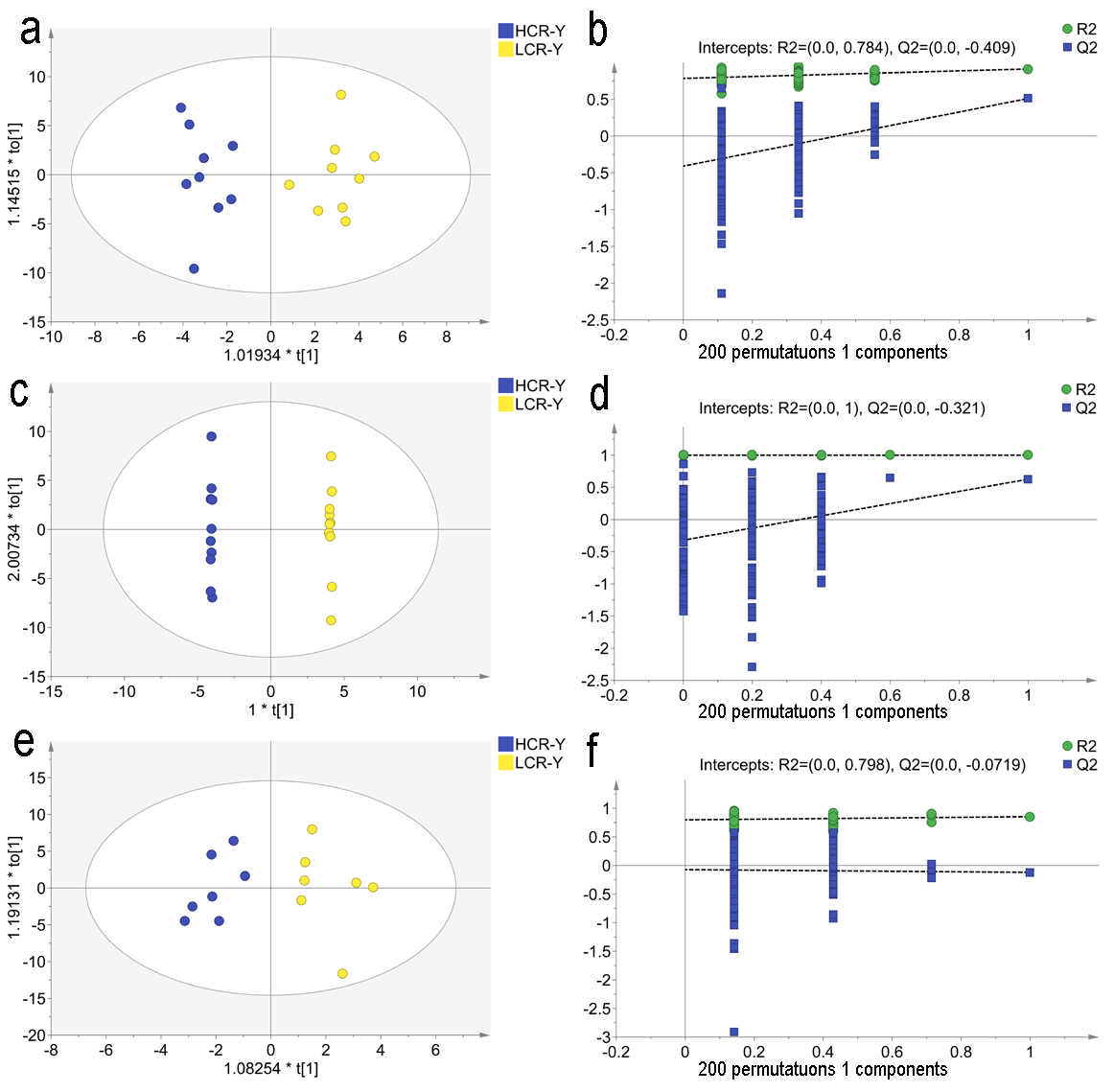

### Supplemental Figure 2

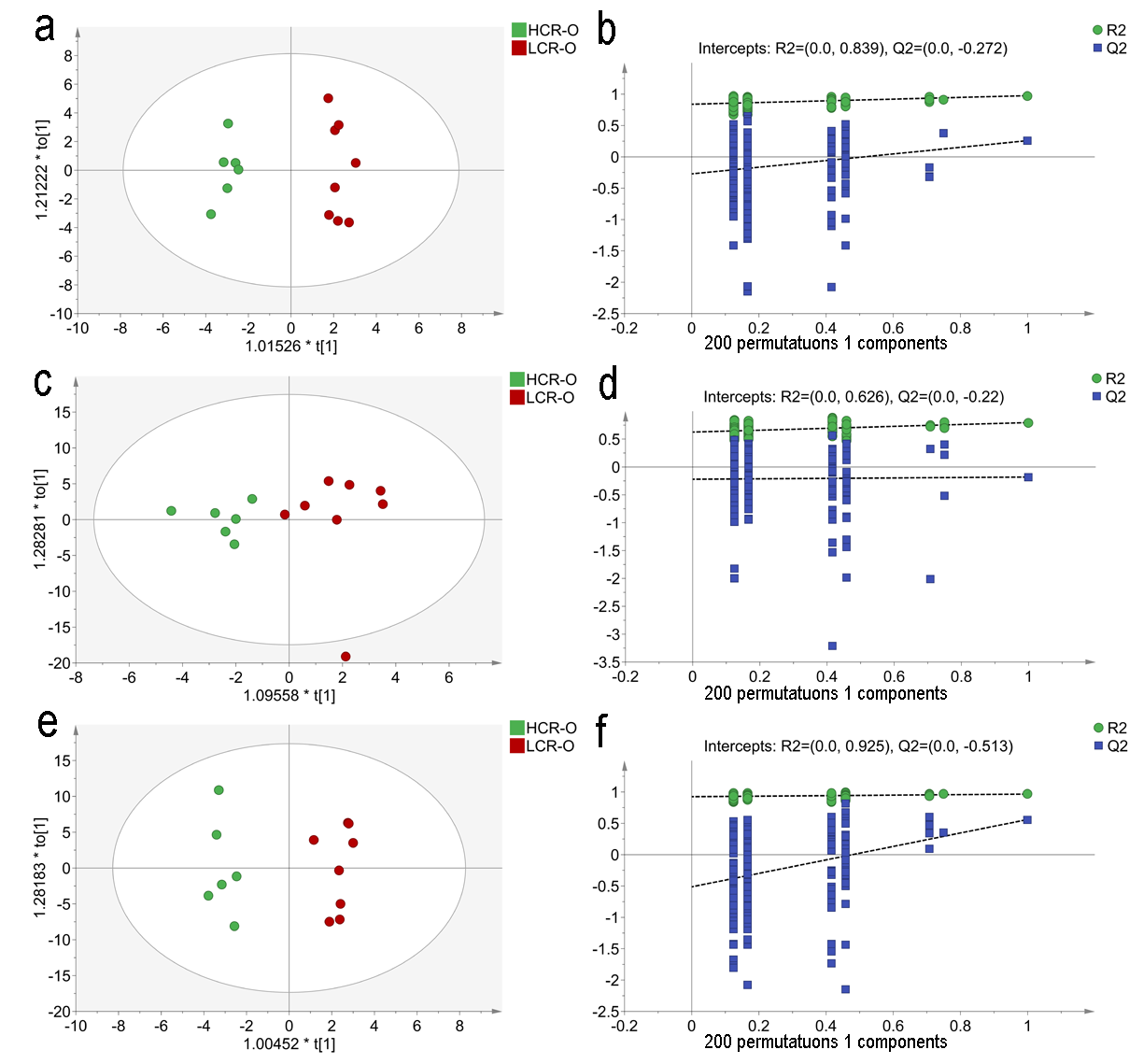

### Supplemental Figure 3

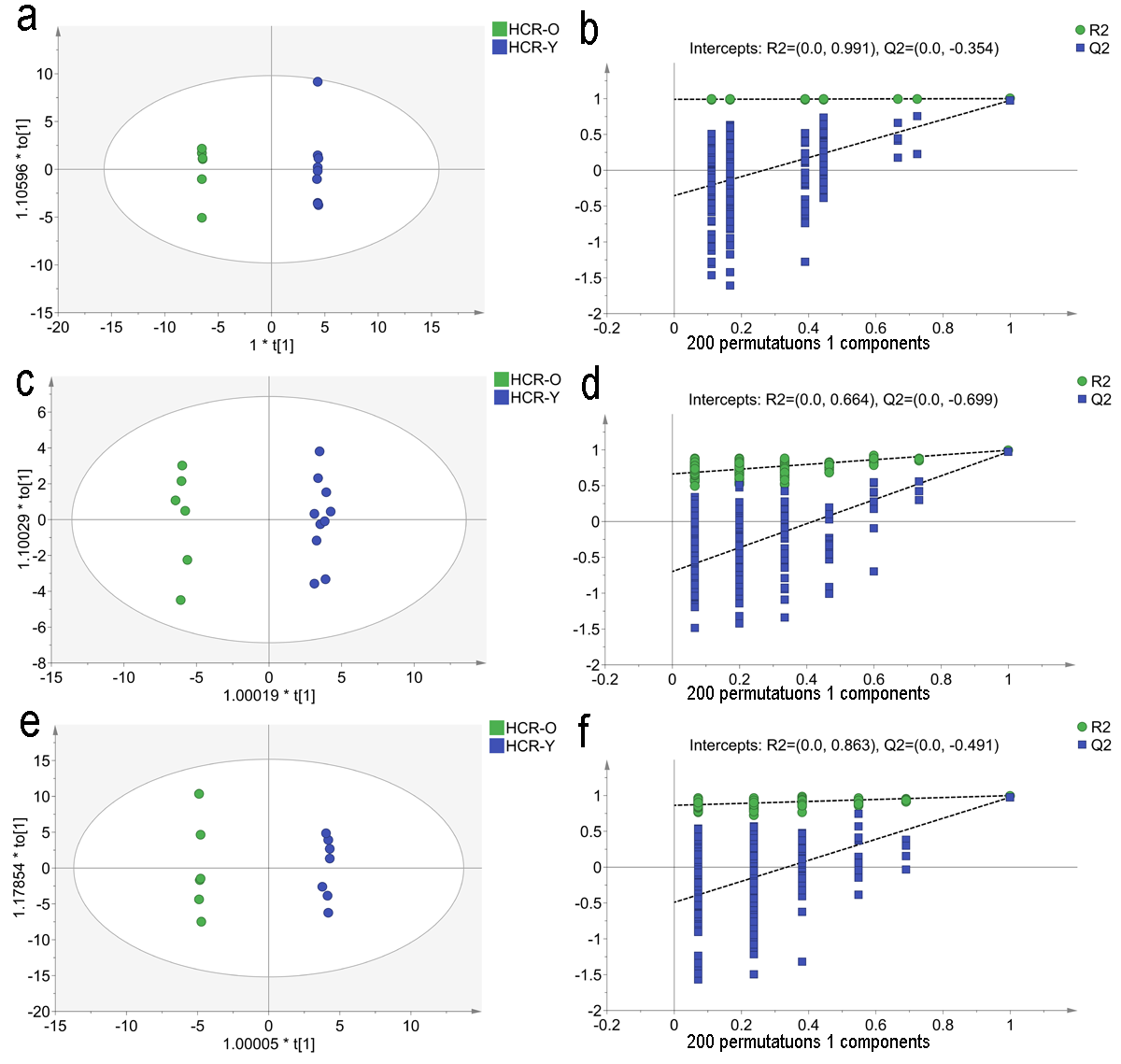

### Supplemental Figure 4

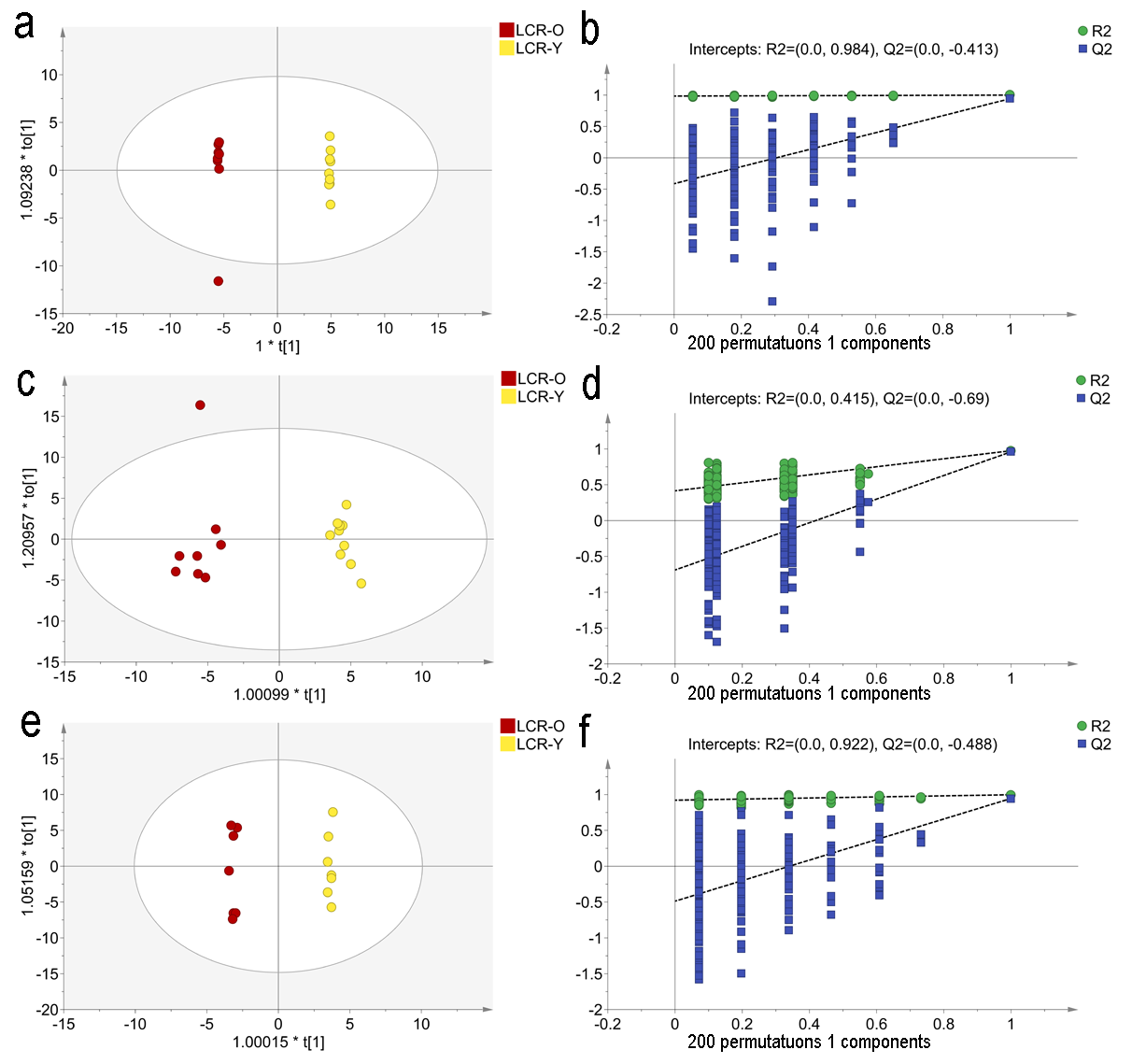
